# Pervasive selection against microRNA target sites in human populations

**DOI:** 10.1101/420646

**Authors:** Andrea Hatlen, Antonio Marco

## Abstract

MicroRNA target sites are often conserved during evolution and purifying selection to maintain such sites is expected. On the other hand, comparative analyses identified a paucity of microRNA target sites in co-expressed transcripts, and novel target sites can potentially be deleterious. We proposed that selection against novel target sites pervasive. The analysis of derived allele frequencies revealed that, when the derived allele is a target site, the proportion of non-target sites is higher than expected, particularly for highly expressed microRNAs. Thus, new alleles generating novel microRNA target sites can be deleterious and selected against. When we analysed ancestral target sites the derived (non-target) allele frequency does not show statistical support for microRNA target allele conservation. We investigated the joint effects of microRNA conservation and expression and found that selection against microRNA target sites depends mostly on the expression level of the microRNA. We identified microRNA target sites with relatively high levels of population differentiation. However, when we analyse separately target sites in which the target allele is ancestral to the population, the proportion of SNPs with high Fst significantly increases. These findings support population differentiation is more likely in target sites that are lost than in the gain of new target sites. Our results indicate that selection against novel microRNA target sites is prevalent and, although individual sites may have a weak selective pressure, the overall effect across untranslated regions is not negligible and should be accounted when studying the evolution of genomic sequences.

## INTRODUCTION

MicroRNAs are small endogenous RNAs that can regulate virtually any type of biological process. Following their discovery in humans this century (Pasquinelli et al. 2000), there are now over 2500 human microRNA precursors annotated in miRBase (Kozomara and Griffiths-Jones 2014), although less than 900 are classified with high confidence. Soon after microRNAs were found in multiple animal species (Lagos-Quintana et al. 2001; Lau et al. 2001; Lee and Ambros 2001), the first target prediction tools became available (Enright et al. 2003; Lewis et al. 2003; Stark et al. 2003). Only in the last few years have these developments permitted the evolutionary analysis of target sites (Farh et al. 2005; Grün et al. 2005; Lewis et al. 2005; Stark et al. 2005; Sood et al. 2006) revealing that many microRNA target sites are highly conserved among species. In contrast, whilst some microRNA families have been conserved for millions of years, their targets appear to differ between species (see for instance (Hui et al. 2013). Indeed, evidence from vertebrates suggests that gains and losses of target sites may be more important than changes in the microRNAs themselves during the evolution of microRNA-based gene regulation, as microRNA genes are usually highly conserved (see Discussion in (Marco 2018a)).

Several studies have found that gene transcripts are depleted of target sites for co-expressed microRNAs (Farh et al. 2005; Stark et al. 2005; Sood et al. 2006). In particular long 3’ UTRs might accumulate microRNA target sites by random mutation, yet they actually have a lower frequency than expected by chance, suggesting that there has been selection against these sequences (Farh et al. 2005). These missing sites have been called ‘anti-targets’ (Bartel and Chen 2004; Farh et al. 2005). Interestingly, target sites for the same microRNAs tend to be conserved in transcripts expressed in neighboring tissues (Stark et al. 2005). These studies have shown that selection against microRNA target sites can be inferred from comparisons among distantly related species. However, the relative impact of selection against microRNA target sites in human populations is not known.

Analysis of human populations has suggested purifying selection was particularly strong at microRNA target sites, even in non-conserved sites (Chen and Rajewsky 2006; Saunders et al. 2007). Negative selection against gaining microRNA target sites has also been described in Yoruban populations, but the pattern was not detected in other populations (Chen and Rajewsky 2006). Correspondingly, we have previously found evidence of selection against microRNA target sites in a study in *Drosophila* populations (Marco 2015). Specifically, we found selection against target sites of maternal microRNAs in maternally deposited transcripts within the egg and early ambryo. More recently it has been shown that this effect is particularly strong for the *mir-309* cluster, whose microRNAs are abundant in the egg and almost absent in the zygote (Zhou et al. 2018). Characterising this type of selection in humans would reveal to which extent it shapes our genomes. However, the strength and prevalence of selection against target sites is human populations is still unknown. Here we investigate single nucleotide polymorphisms at human microRNA target sites and evaluated the impact of selection against target sites. To do so, we consider pairs of segregating alleles in which one of the allele as a target site and the other was not. Then we compare the allele frequency distributions with that of estimated background distributions to quantify the strength of selection for or against microRNA target sites.

## RESULTS

### Bias towards microRNA non-target alleles in human populations

In order to investigate the selective pressures on microRNA target sites in human populations, we first mapped human single-nucleotide polymorphisms (SNPs) to putative canonical microRNA target sites such that one allele is a target site and the alternative allele is not a target site (see Methods for details). The non-target allele in this pair is called a ‘near-target’ (Marco 2018b). We first compared the derived allele frequency (DAF) distribution of target sites for highly expressed microRNAs with a background (expected) distribution obtained by conducting the same analysis on the reverse complement sequences of 3’UTRs (see Methods). We considered those SNPs for which the derived allele is the target allele. If the overall selective pressure is to fix new target sites (positive selection favouring new interactions), DAF values should be the higher than expected. On the other hand, if there is selection against microRNA target sites, the DAF distribution should be skewed towards smaller values (Figure 1). We observe an overall excess of low frequency derived alleles (Figure 2A; p=0.009, one-tailed Kolmogorov-Smirnov test).

**Figure 1.**
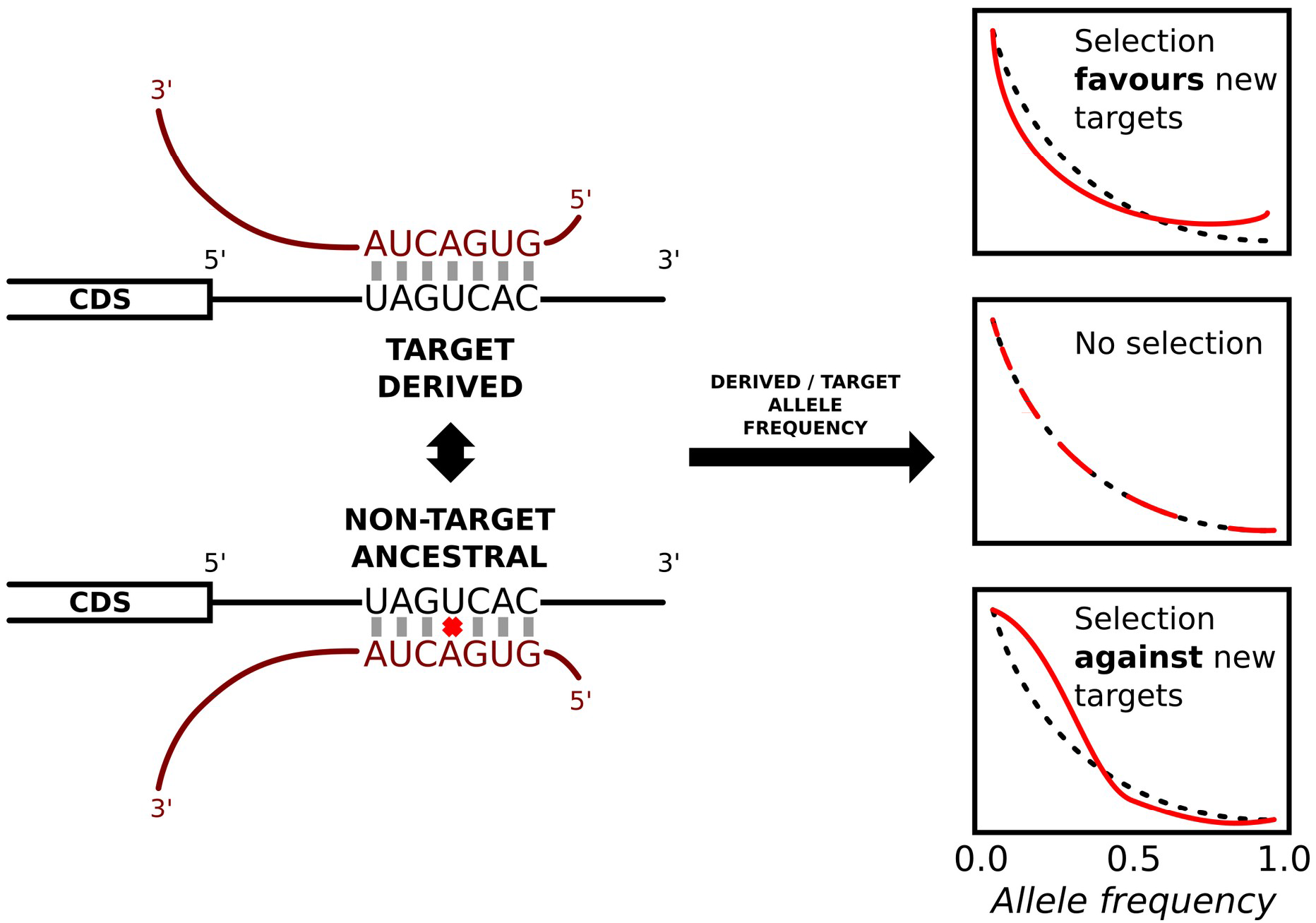
Analysis of derived allele frequencies at target sites. Pairs of alleles where one allele is a target site and the other is not are identified, and only those where the non-target allele was the ancestral state were further considered. The derived (not ancestral) allele frequency distribution will be skewed towards the target allele (right) if selection favours the emergence of new microRNA target sites, or to the non-target allele (left) if selection acts against new targets.

**Figure 2.**
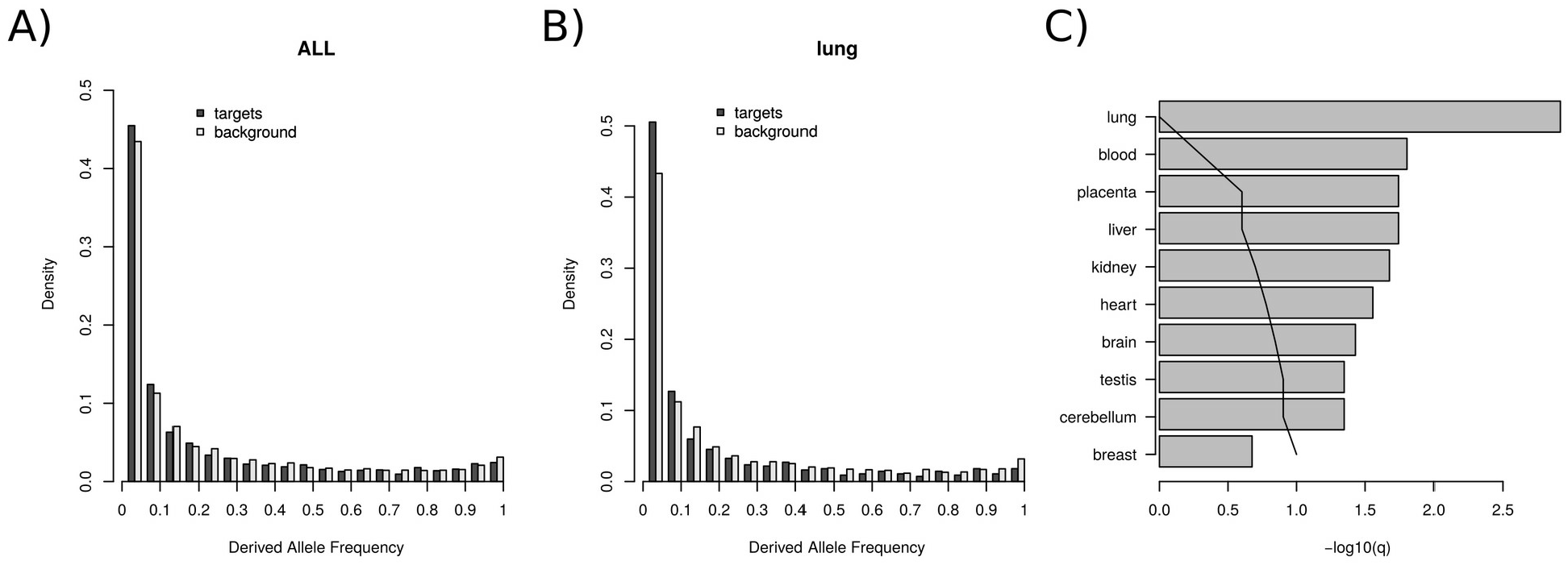
Derived allele frequencies (DAF) at microRNA target sites. (A) Derive allele frequency distribution of microRNA target sites, where the derived allele is a target site, compared to a background distribution for the whole dataset. (B) As in A but for interactions between coexpressed microRNA/transcript pairs in lung tissue. (C) Significance levels for differences in the DAF distribution for ten tissues expressed as −log10[q-value] (see Methods). The vertical line indicates the threshold for one expected false positive.

It is expected that a selective pressure on co-expressed microRNA/mRNA pairs will be tissuespecific. Therefore, we compared the DAF distributions as above for ten specific tissues, considering only interactions in which the microRNA and the potential target are co-expressed. For instance, for lung co-expressed microRNA/transcripts we found a DAF distribution significantly skewed towards the non-target allele, indicating selective pressures against the target allele (one-tailed Kolmogorov-Smirnov test, p=0.00012; Figure 2B). For all tissues analysed we observe a shift in the derived allele frequencies towards the non-target allele (Supplementary Figure 1), all but breast within a 10% False Discovery Rate (less than one expected false positive; Figure 2C, Supplementary Table 1). The poor statistical signal in breast tissue can be easily explained by a very small sample size (42 SNPs in total).

Genes can have alternative polyadenylation in different tissues, affecting microRNA target sites. However, in the context of microRNAs, this has been explored mostly in cell lines (Nam et al. 2014). We found alternative polyadenilation information for two of the tissues here investigated: blood and kidney (Müller et al. 2014). When we consider target sites whose 3’UTR has been experimentally detected in those tissues, the shift to the non-target allele is significant (Supplementary Table 1; Supplementary Figure 1).

Alternatively, we can compare the allele frequency distributions between target sites for highly expressed microRNAs versus target sites for non-expressed (non-detected) microRNAs in a given tissue. To do so we identified microRNAs with no read counts detected across multiple expression experiments (see Methods) and used their target sites as our background (expected) frequencies. Consistently with the results above, we found that for most analysed tissues the DAF distribution when the ancestral allele is a non-target is skewed to the non-target allele (Supplementary Figure 2, Supplementary Table 2). In summary, the distribution of allele frequencies shows evidence of selection against microRNA target sites in human populations. Again, the results are consistent when taking into account alternative polyadenylation in blood and kidney (Supplementary Table 2; Supplementary Figure 2).

Wobble pairing has been also observed in microRNA/transcript interactions. Under our hypothesis, selection against target sites will be weaker at sites in which the non-target allele is a potential target with a wobble pairing (GU). Hence, we partitioned the dataset of targets in those whose near-target allele is a wobble paired position and those which are not. We observed that for near-target-wobble the shift to the non-target allele is not significant (0.682) whilst for the non-wobble there was a significant shift (0.009, Supplementary Figure 3). These results furthers strengths the evidence for selection against microRNA target sites.

### The effect of microRNA expression levels and evolutionary conservation

We next considered the potential impact of microRNA conservation. On the one hand, evolutionarily conserved microRNAs may have a weaker effect on selection against microRNA targets, as partly deleterious target alleles may have been cleared from the population. On the other hand, evolutionarily conserved microRNAs tend to be highly expressed, and therefore it is expected that the selective pressure to avoid targets for such microRNAs should be stronger. In other words, expression and conservation are not independent to each other. We therefore included both factors in the analysis: the level of expression of the microRNA and a measure of the phylogenetic conservation of the microRNA sequence.

We analysed cases where the derived allele is a target site (as in the previous section); testing whether frequency spectra were different across different levels of microRNA conservation (human-primate specific, conserved in mammals, and conserved in animals) and different levels of expression (low, mid and high as described in Methods). To evaluate the joint effect of conservation and expression, we build a linear model of ranked independent variables with interactions (this is equivalent to the Scheirer-Ray-Hare test ((Sokal and Rohlf 1995), pp.445; see Methods). The interaction term was not statistically relevant in any of the models (Supplementary Table 3). In general, from the fitted model it is evident that expression has a significant impact in the microRNA selective avoidance whilst conservation has no detectable impact, once the expression level has been taken into account (Figure 3, Supplementary Table 3).

**Figure 3.**
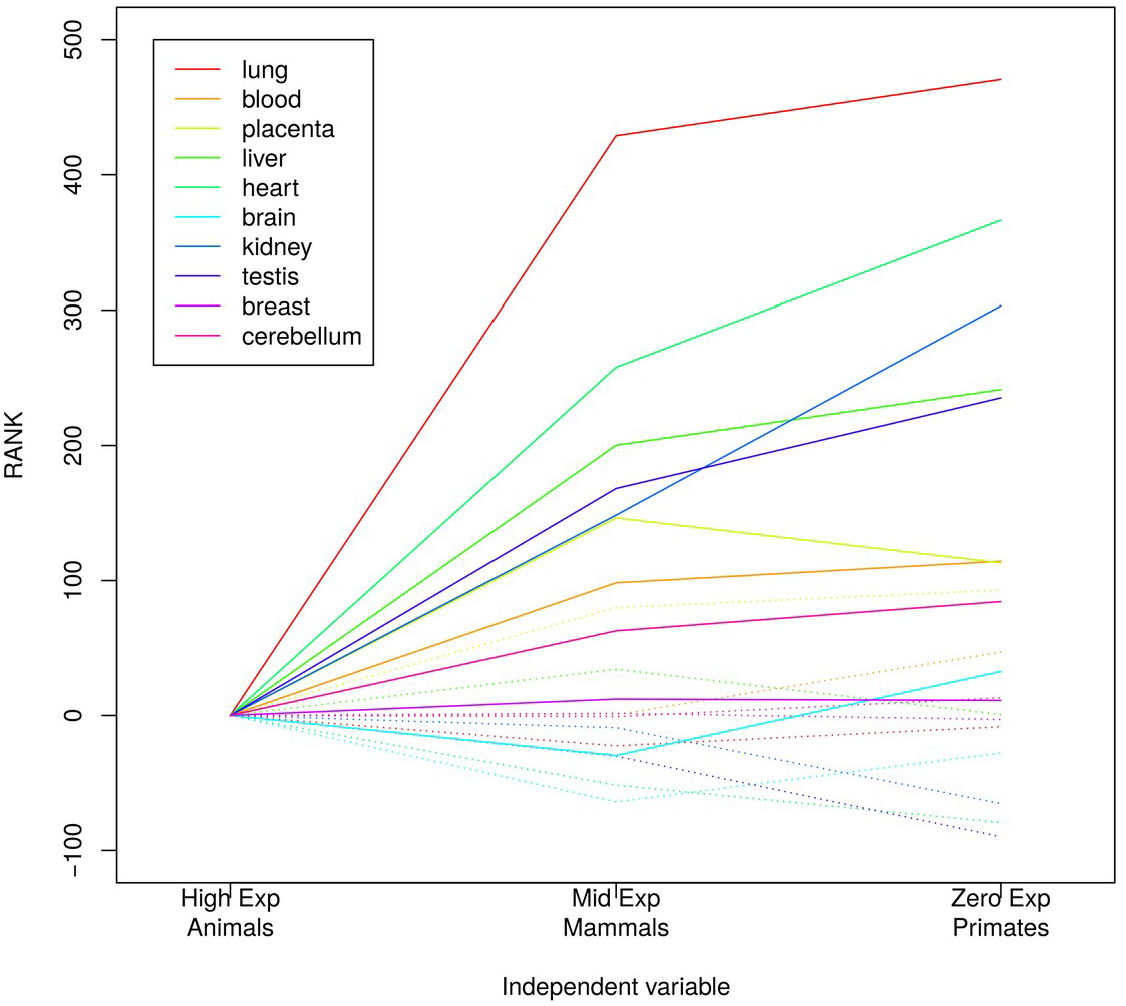
The effect of microRNA conservation and expression level on the frequency of target alleles. Rank differences in our linear model (see Methods) explained by expression (solid lines) and conservation (dashed lines) levels for ten different tissues. Highly expressed and conserved in animals microRNAs were taken as the intercept of the model (rank zero).

More specifically, in Figure 2 the contribution of expression and conservation to the ranks in the linear model are plotted for all ten tissues. The smaller the rank, the greater the shift to the ancestral non-target allele in a comparison of derived allele frequencies. Overall, we observe that as the expression level increases the rank decreases, whilst the rank is similar for all three conservation levels (Figure 2). The effect is particularly clear in lung, liver and kidney. For brain ans breast we did not find such an association (Figure 2; Supplementary Table 3). From these analyses we concluded that expression level, rather than microRNA evolutionary conservation, determines the selective pressure against microRNA target sites.

### Population differentiation at target sites

If a microRNA target site is under selective constraints, we should expect differentiation among populations at these sites to be relatively low. To investigate this prediction, we grouped SNPs at target sites for highly expressed microRNAs according to their Fst and compared the relative frequency of these SNPs compared to the background (see Methods). However, in Figure 3A we observed an enrichment in high Fst values. To further explore the relative contribution of microRNA target gains and losses during population differentiation we split the dataset in two groups, depending on whether the target allele was ancestral or derived. Strikingly, polymorphic microRNA target sites where the ancestral allele is the target show high levels of population differentiation (Figure 3A, red line). In contrast, for novel microRNA target sites there is a deficit of high Fst SNPs compared to the background expectations (Figure 4A, blue line). When we repeated the analysis for moderately expressed microRNA we found a similar pattern (Figure 4B). This may reflect that positive selection driving the generation of novel microRNA target sites is negligible. Also, evolution by microRNA target site loss seems important in human populations. In Table 1 we show SNPs in microRNA target sites with a Fst greater than 0.6. As suggested in Figure 4A and 4B, in a majority of target sites with a high degree of population differentiation the target allele was ancestral (21 out of 35), many being probably target losses in out-of-Africa populations (12 out of 21). Predicted target sites for microRNAs with no detectable expression level did not show any significant level of population differentiation (Figure 4C).

**Figure 4.**
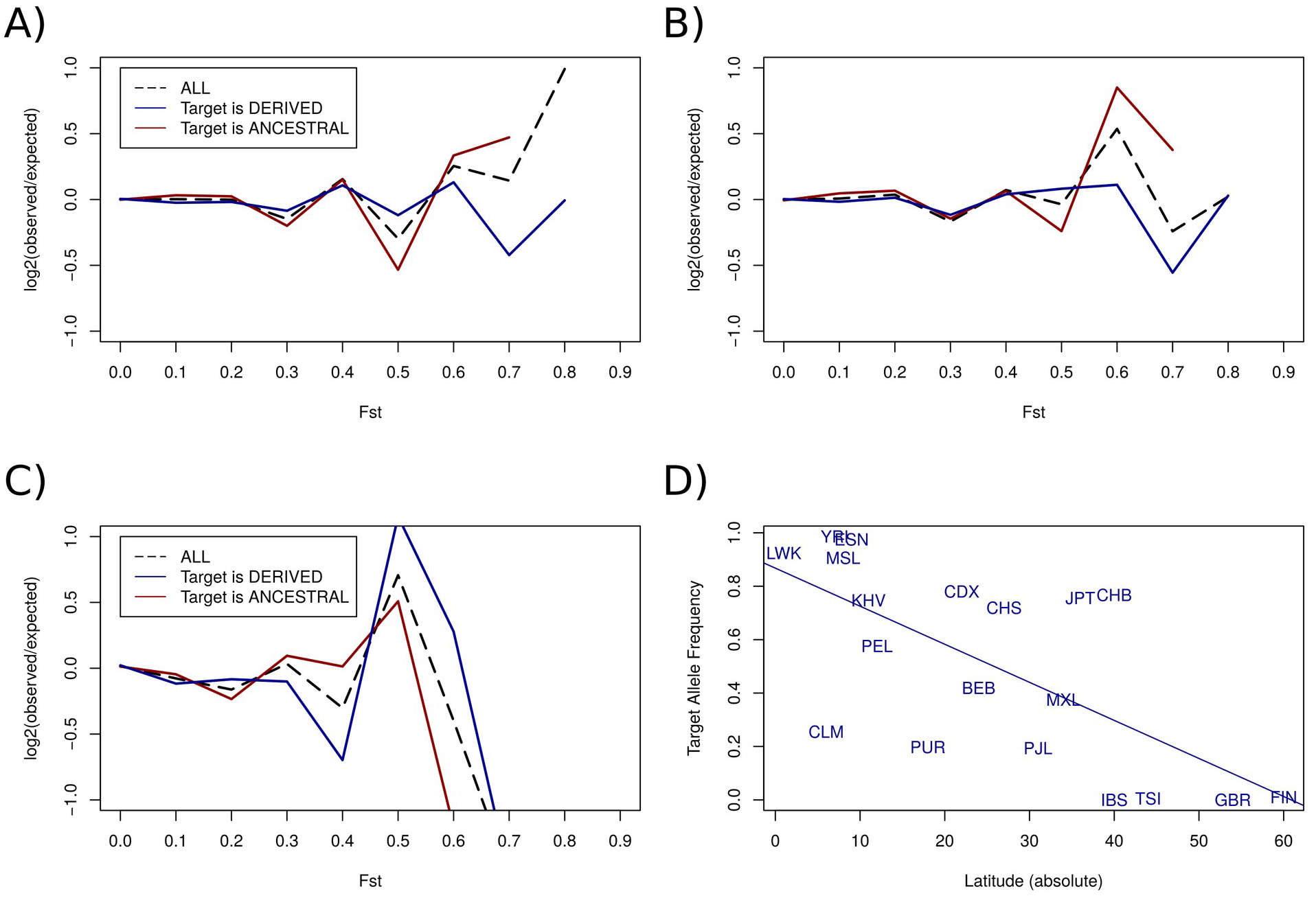
Fst enrichment in polymorphic microRNA target sites. Comparisons of the Fst values for SNPs within microRNA target sites with control sites (background sites, see Methods). For each range of Fst values, the proportion of sites in that range was calculated for the microRNA target sites (*P_t_*) and for the background sites (*P_b_*). The value plotted is the log(*P_t_*/*P_b_*). (A) All targets sites for highly expressed microRNAs (black, dashed line); target sites where the target allele is ancestral (red) and targets sites where the target site is not ancestral (blue). (B and C) As in (A) but for moderately expressed and zero expressed microRNAs respectivelly. (D) Regression between absolute latitude in samples human populations and the target allele frequency associated to SNP rs2470102.

**Table 1.** Target allele frequencies at polymorphic sites with a high Fst values.

| miRname<sup>1</sup> | Gene | snpName | target<sup>2</sup> | Target Allele Frequencies |  |  |  |  |  |  |
| --- | --- | --- | --- | --- | --- | --- | --- | --- | --- | --- |
|  |  |  |  | EAS<sup>3</sup> | AMR | AFR | EUR | SAS | ALL | Fst |
| miR-92a-3p | PTK6 | rs186332 | d | 0.964 | 0.873 | 0.047 | 0.928 | 0.593 | 0.630 | 0.849 |
| miR-518d-5p | MTAP | rs7868374 | a | 0.008 | 0.048 | 0.684 | 0.005 | 0.118 | 0.213 | 0.824 |
| let-7a-5p | MTAP | rs7875199 | a | 0.054 | 0.058 | 0.796 | 0.007 | 0.129 | 0.256 | 0.800 |
| miR-202-5p | ATP1A1 | rs1885802 | a | 0.043 | 0.127 | 0.811 | 0.034 | 0.045 | 0.256 | 0.791 |
| miR-1180-3p | MYEF2 | rs2470102 | a | 0.753 | 0.339 | 0.925 | 0.006 | 0.272 | 0.497 | 0.773 |
| miR-24-3p | SCN2B | rs624328 | d | 0.941 | 0.891 | 0.252 | 0.951 | 0.967 | 0.760 | 0.767 |
| miR-21-3p | C4orf46 | rs11544037 | a | 0.025 | 0.262 | 0.854 | 0.130 | 0.169 | 0.326 | 0.745 |
| miR-130a-3p | SLC30A9 | rs12511999 | a | 0.049 | 0.236 | 0.913 | 0.250 | 0.222 | 0.377 | 0.730 |
| miR-513a-3p | TCERG1 | rs3822506 | d | 0.718 | 0.117 | 0.033 | 0.092 | 0.220 | 0.231 | 0.722 |
| miR-1296-5p | BCL7C | rs11864054 | a | 0.088 | 0.455 | 0.976 | 0.618 | 0.841 | 0.628 | 0.714 |
| miR-192-5p | C12orf65 | rs1533703 | a | 0.998 | 0.726 | 0.149 | 0.769 | 0.795 | 0.651 | 0.707 |
| miR-221-5p | FZR1 | rs10155 | d | 0.798 | 0.473 | 0.015 | 0.294 | 0.302 | 0.348 | 0.692 |
| miR-7-5p | ENAM | rs7665492 | a | 0.023 | 0.133 | 0.744 | 0.058 | 0.055 | 0.242 | 0.685 |
| miR-514a-3p | EXOC5 | rs3742577 | a | 0.114 | 0.125 | 0.845 | 0.119 | 0.121 | 0.311 | 0.683 |
| miR-769-5p | ARIH1 | rs11072379 | a | 0.235 | 0.212 | 0.894 | 0.095 | 0.097 | 0.351 | 0.674 |
| miR-769-5p | MPHOSPH9 | rs1727314 | a | 0.977 | 0.726 | 0.142 | 0.768 | 0.741 | 0.634 | 0.668 |
| miR-381-3p | PLXNA4 | rs6968754 | a | 0.166 | 0.445 | 0.920 | 0.285 | 0.361 | 0.466 | 0.658 |
| miR-520d-3p | UBE2Q1 | rs11265634 | a | 0.098 | 0.221 | 0.870 | 0.186 | 0.199 | 0.356 | 0.649 |
| miR-513c-5p | CYB5R4 | rs6912739 | d | 1.000 | 0.932 | 0.381 | 0.954 | 0.994 | 0.818 | 0.646 |
| miR-335-3p | PLCB2 | rs4257181 | a | 0.446 | 0.899 | 0.987 | 0.991 | 0.911 | 0.853 | 0.643 |
| miR-19b-3p | TRNT1 | rs60884103 | d | 0.699 | 0.242 | 0.054 | 0.024 | 0.360 | 0.264 | 0.643 |
| miR-22-3p | OPLAH | rs28475718 | d | 1.000 | 0.957 | 0.437 | 0.989 | 0.969 | 0.838 | 0.641 |
| miR-335-3p | ANGEL2 | rs41277158 | a | 0.997 | 0.950 | 0.412 | 0.963 | 0.949 | 0.821 | 0.641 |
| miR-103a-3p | HEXA | rs11629508 | d | 0.774 | 0.839 | 0.179 | 0.946 | 0.943 | 0.694 | 0.631 |
| miR-34a-5p | CDPF1 | rs1053332 | a | 0.001 | 0.169 | 0.746 | 0.194 | 0.082 | 0.276 | 0.631 |
| miR-144-3p | CFAP61 | rs1410937 | a | 1.000 | 0.954 | 0.412 | 1.000 | 1.000 | 0.839 | 0.626 |
| miR-144-5p | RSU1 | rs6977 | d | 0.895 | 0.921 | 0.275 | 0.982 | 0.932 | 0.760 | 0.625 |
| miR-223-3p | PCDH15 | rs11003862 | d | 0.055 | 0.591 | 0.731 | 0.764 | 0.440 | 0.526 | 0.625 |
| miR-145-5p | C3orf85 | rs56027044 | d | 0.164 | 0.392 | 0.879 | 0.590 | 0.441 | 0.524 | 0.622 |
| miR-509-5p | PKDREJ | rs6007729 | a | 1.000 | 0.937 | 0.421 | 0.963 | 0.964 | 0.825 | 0.621 |
| miR-197-3p | GP2 | rs12444232 | d | 0.682 | 0.157 | 0.032 | 0.047 | 0.190 | 0.214 | 0.616 |
| miR-200b-3p | PCDH15 | rs11003861 | a | 0.810 | 0.339 | 0.048 | 0.179 | 0.454 | 0.347 | 0.612 |
| miR-124-3p | CPSF4 | rs1043466 | d | 0.660 | 0.716 | 0.055 | 0.833 | 0.618 | 0.535 | 0.608 |
| miR-381-3p | INAFM2 | rs2289333 | d | 0.554 | 0.308 | 0.007 | 0.090 | 0.153 | 0.204 | 0.607 |
| miR-377-3p | OGDHL | rs6816 | a | 0.105 | 0.529 | 0.933 | 0.553 | 0.582 | 0.566 | 0.607 |

<sup>1</sup>Representative microRNA of the family; <sup>2</sup>This column indicates if the target allele is ancestral ('a') or derived ('d'); <sup>3</sup>Target allele frequency for the five super-populations analysed (see Methods for definitions).

One of the SNPs in Table 1 has been recently associated with skin colour variation in Indian populations: rs2470102 (Sarkar and Nandineni 2018). Interestingly, this is in a predicted a target site for miR-1180-3p in MYEF2. This gene has been associated with skin colour as well, although the functional relationship is not clear (Mishra et al. 2017). We investigated a potential relationship between absolute latitude (distance to the Ecuador) and the frequency of the target allele and we found a strong association (Figure 4D; p=0.0015, R-squared adjusted = 0.4223). Given that the microRNA itself originated in primates (Arcila et al. 2014), and that the target allele is ancestral and conserved, the emergence of a new microRNA may have imposed a selective pressure to loss the target site in populations with less exposure to UV light. However interesting, these association remains speculative at the moment, and demographic differences between populations may also have an impact in this observed association.

These results further suggests that loss rather than conservation of targets may be more frequent in population dynamics. Thus, we revisited the analysis of derived allele frequencies in Figure 2 and considered this time target sites where the ancestral allele was a target site. If there is a detectable selective pressure to maintain target sites we will expect the derived allele frequency to be biased towards the target (ancestral) allele. In Figure 5 we plot the bias (as the D-statistic in a Kolmogorov-Smirnov test) and the significance (as the log10 of the p-value) and we observe that in this set the bias was not significant (blue dots). On the contrary, if we compare these results with those obtained for non-targets as ancestral sequences we observe a clear and significant biased towards the non-target allele (Figure 5).

**Figure 5.**
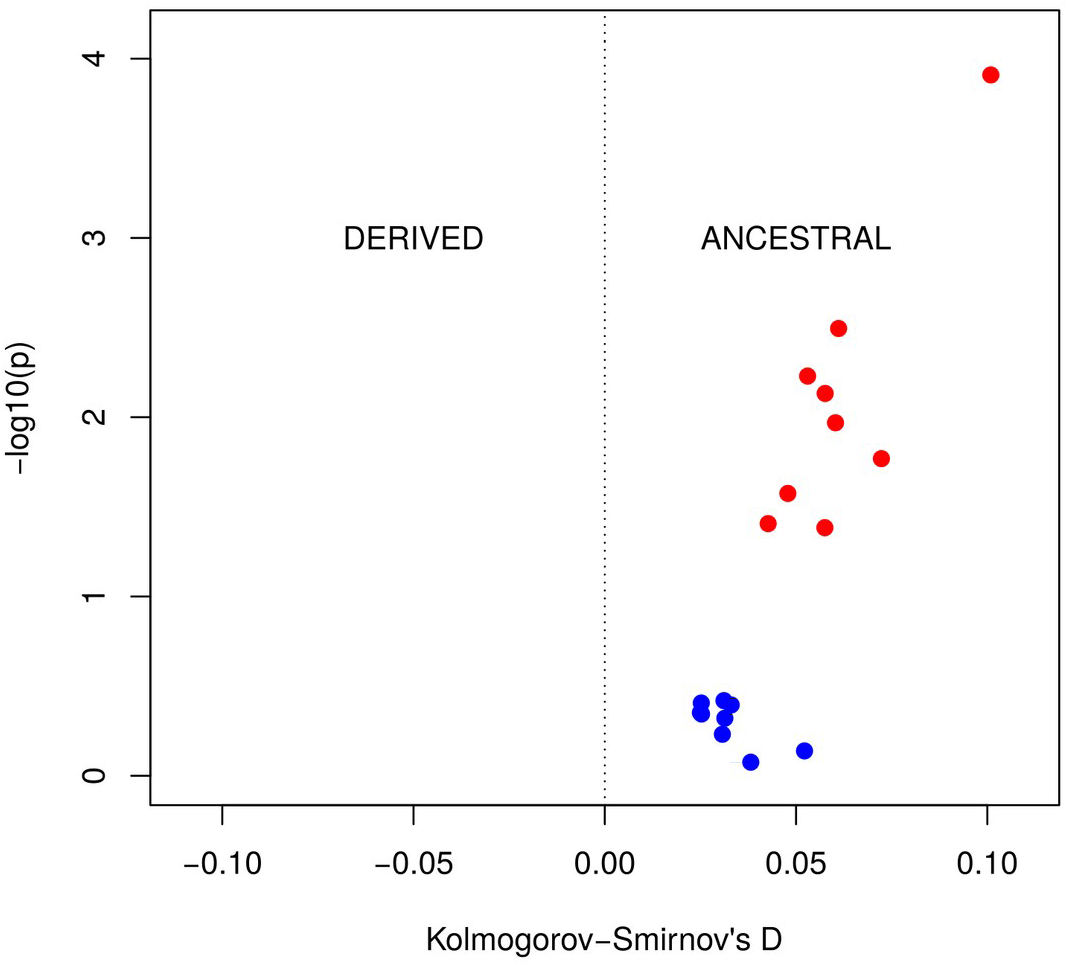
Allele preference for ancestral targets and non-targets. The shift of the distribution of derived allele frequencies is measured as the D statistic form the Kolmogorov-Smirnov test (x-axis). When the distribution is biased towards the derived allele the D statistic is negative (-D^-^), and if the bias is to the ancestral allele the D value is D^+^. The y-axis plots the −log10 of the p-value obtained from the corresponding one-tailed KS test. For each tissue the graph represents the DAF bias when the ancestral allele is a target (blue circles) and when it is a non-target (red circles).

In one case we found a microRNA with a SNP within its seed sequence (the region that determines the targeting property of the microRNA), which shows some evidence of population differentiation (rs7210937; Fst = 0.3314). In this case, the Fst value between African and European populations is remarkably high (Fst = 0.6129; Nei’s estimate (Nei 1986; Bhatia et al. 2013)). In European populations 92.5% of sequenced individuals present the ancestral form of miR-1269b, whilst in African population, the derived version is more frequent (59.8%). As a shift in the seed sequence may have an impact on the evolution of 3’UTRs, we further studied target sites whose ancestral form is a target for the derived miR-1269 microRNA (344 in total). Then we compared the frequency of the target site allele between European and African populations. We found that in African populations the frequency of target alleles is lower than in European populations at these sites (Wilcoxon non-parametric test for paired samples; p=0.021) whilst for the ancestral miR-1269b we did not find any significant difference. These results suggest that a shift in the allele frequencies affecting the seed sequence of a microRNA can have an effect on the allele frequencies at the novel target sites, specifically toward the non-target allele.

## DISCUSSION

The study of allele frequencies has been extensively used to detect selective pressures in human populations (Chen and Rajewsky 2006; Barreiro et al. 2008; Enard et al. 2014). Here we show that the patterns of allele frequencies at 3’UTRs show evidence of selection against most microRNA target sites. First, the allele frequencies at target sites are biased towards the non-target allele when the derived allele is a target sequence. These effects are strongest in the cases where the corresponding microRNAs are highly expressed, suggesting that interaction between the microRNA and the target is a key is the source of selection against target sequences. The microRNAs that have been conserved over longer periods of vertebrate evolution did not impose detectably greater selection against their target sequences, once the effect of expression levels had been taken into account. On the other hand we failed to detect any noticeable effect when the target sites was ancestral, and therefore we did not find statistical signal supporting selection to maintain existing microRNA target sites. We did not believe that means that there is no purifying selection to keep functional sites, but that this is comparably small compare to selection to avoid new, potentially deleterious, microRNA target sites. As a matter of fact, novel microRNA target sites has been identified in various studies as disease causing mutations (Abelson et al. 2005; Dusl et al. 2015). The most popular microRNA target prediction programs rely on target site conservation to reduce the number of false positives (Agarwal et al. 2015) and/or do not provide a stand-alone version to run on custom datasets (Paraskevopoulou et al. 2013). Therefore, we used a naive microRNA target prediction method that reports canonical targets and near-target sites (Marco 2018b). That allowed us to study pairs of alleles segregating at target sites without any other constraint. On the other hand, we would expect a high number of false positive in target predictions (reviewed in (Alexiou et al. 2009)). Remarkably, we found a significant pattern of selection against microRNA target sites. This reinforces our initial hypothesis and suggest that, if we would be able to restrict the analysis to bona fide target sites, the signal might be stronger. One possibility is to evaluate experimentally validated target sites. However, these experiments are based on reference genomes, so segregating target sites whose target allele is not in the reference genome will be lost from the analysis. The way forward may be to perform high-thoughtput microRNA target experiments, like HITS-CLIP (Chi et al. 2009), in cells derived from different populations. The continuous drop in the costs of sequencing and high-throughput experiments may allow this in the near future. Indeed, high-throughput experimental evaluation of segregating alleles at regulatory motifs (transcription factor binding sites, RNA binding sites, etcetera) is a promising area of research which will help us to move from a typological (reference genome) to a population view of gene regulation.

Another way to study the effect of selection in populations is to evaluate the population differentiation (Coop et al. 2009; Li et al. 2012). We found that, at microRNA target sites in general, there is an enrichment in high population differentiation. This result was observed by by Li et al. (Li et al. 2012). However, we found that this trend only holds for those sites at which the ancestral allele was a microRNA target site and the derived allele is a non-target. The loss of a microRNA target (as in the examples reported in a previous work (Li et al. 2012)) may be relatively frequent. It follows that the non-target allele might be neutral or even advantageous in some of these cases. It is noticeable that in the examples in which the derived allele reaches a high frequency, that occurs in the non-African populations, which is the pattern that would be expected if a neutral derived allele spread by genetic drift during founder events. Loss of target sites could also have advantageous effects, though the complex interactions that occur in regulatory networks. For example, it has been proposed that in the human lineage the loss of microRNA target sites contributed to an increase in the expression levels of some genes (Gardner and Vinther 2008). We also recently reported the loss of multiple target sites in Out-of-African populations (Helmy et al. 2019). Our work suggest that this loss of targets may be continuing now in human populations. Selection in favour of new target sites appears to be rarer: we found a strong signal of purifying selection against novel microRNA target sites.

It is worth mentioning that our result as well as other works are based on precomputed Fst values that may be affected by systematic biases in allele imputation. As more genome sequences of high quality are released, and new Fst estimations become available, it will be important to re-evaluate the impact of population differentiation in microRNA target sites. In general, our results also depend on the ability to predict microRNA target sites from primary sequences. As the number of polymorphic microRNA target sites experimentally validated is very limited, at the moment our approach is the only possible. However, it’ll be important to re-evaluate our results in due course as more datasets become available. Also, our statistical analysis relies on two types of null models: microRNAs with not detected expression, and the analysis of the reverse complement sequence of 3’UTRs. It is possible that microRNAs expressed in only a few cells are wrongly attributed to the ‘null’ class. Likewise, the reverse complement of 3’UTR could also be potentially transcribed (antisense transcripts) and, hence, wrongly selected as ‘null’ class. In any case, the use of complementary null models leading to similar results supports our hypothesis, and indicates that the use of better annotated datasets in the future may even increase the statistical evidence in favour of selection against microRNA target sites.

Our results suggest that new microRNA target site generating mutations (the derive allele is the target) are selected against. This is a case of the classic selection-mutation balance in which the mutations are deleterious. Selection against deleterious mutations has been extensively studied in population genetics (reviewed in (Charlesworth 2012)). For instance, strong purifying selection produces a phenomenon called background selection, in which loci linked to the selected site experience a reduction in their effective population size (Charlesworth et al. 1993). That is, purifying selection reduces the influence of selection at linked sites. For weakly selected sites, a similar process has been described: weak selection Hill-Robertson interference (wsHR (Hill and Robertson 1966; McVean and Charlesworth 2000)). Under wsHR, multiple alleles are under a weak selective force, very close together so that recombination is small or negligible between sites, interfering with other selective pressures in the area. Multiple weakly deleterious mutation at transcription factor binding sites have been reported indeed (Abecasis et al. 2012). We believe that this is the case for the selection against microRNA target sites here described: weak selection against multiple target/near-target sites will shape the evolutionary landscape of the entire untranslated region.

Our study suggest that the mutation rate in humans may be high enough to produce a significant selective pressure against novel microRNA target sites. New target sites will emerge at a significant rate because many mutations can potentially introduce a new site for one of the many microRNAs. More specifically, there are about 2,000 microRNA families described in TargetScan (see Methods), defined by 7-nucleotide seed sequences. Assuming that 3’UTRs are composed of non-overlapping 7-mers (a simplifying yet conservative assumption) the expected number of near-target sites per kilobase (kb) is about 75. With a mutation rate of 2.5 10^-5^ per kb (Nachman and Crowell 2000) and a total length of the genome that encode 3’UTRs of 34Mb, it can be shown that there will be on average one novel microRNA target site on a 3’UTR per genome per generation. That is, one potential deleterious microRNA target site per person per generation.

It is expected that other regulatory motifs influence the evolution of 3’UTRs. For instance, Savisaar et al (Savisaar and Hurst 2017) have described selection against RNA-binding motifs. The selective avoidance of transcription factor binding sites (Hahn et al. 2003; Babbitt 2010) and of mRNA/ncRNA regulatory interactions in bacteria (Umu et al. 2016) have been also described. These works are based on comparative genomics between different species and not on variation within populations. However, some theoretical models take into account the selective pressure against regulatory motifs (Berg et al. 2004; Stewart et al. 2012). It is likely that on top of all the selective forces that are usually taken into account, there is a layer of selection against weakly deleterious regulatory motifs that will be influencing the evolution of the genome. In conclusion, selection against microRNA target sites in prevalent in human populations, and it may constrain other selective forces in post-transcriptional regulatory regions.

## METHODS

MicroRNA mature sequences were downloaded from miRBase v.22, considering only sequences annotated with ‘high-confidence’ (Kozomara et al. 2018). MicroRNA target and near-target sites were predicted with seedVicious (v.1.1 (Marco 2018b)]) against 3’UTR as annotated in Ensembl version 96 (Kinsella et al. 2011) for the human genome assembly hg38. Single nucleotide polymorphisms for the 1000 Genomes project (1000 Genomes Project Consortium et al. 2015) were retrieved from dbSNP (build 151) (Sherry et al. 2001) and mapped to our target predictions. Ensembl sequences and polymorphism data were downloaded using the BiomaRt R package (Durinck et al. 2009). By mapping SNPs to targets and to near-targets (as described in (Marco 2018b)) we are able to identify pairs of alleles in which only one of the allele is a target site, so allele frequencies can be computed as target allele frequencies (Hatlen and Marco 2019). When plotting allele frequencies (Figure 2) we only considered segregating alleles in which the minor allele is present in at least 1% of the sampled population, as reported dbSNP (see (Hatlen et al. 2019) for details). The ancestral allele status was also retrieved from the pre-computed values available from dbSNP. To compute the background (randomly expected) allele distributions we repeated the process but finding targets in the reverse complement strand of the 3’UTR, to control for sequence length and composition. In the analysis of allele frequencies we used a second microRNA target prediction program based on a different principle, miRanda (Enright et al. 2003), which takes into account the binding energy of the RNA:RNA duplex, and we use the default parameters. About 25% of polymorphic canonical target sites were also predicted as targets by miRanda.

Expression information for microRNAs was obtained our own database PopTargs (https://poptargs.essex.ac.uk; Hatlen and Marco 2019). In summary, high-throughput experiments from Meunier et al. (Meunier et al. 2013) and miRMine (Panwar et al. 2017) were downloaded, reads were mapped to miRBase mature microRNA sequences and the average reads per million for each microRNAs was consider to measure the expression level in a tissue. Coding genes were considered to be expressed in a given tissue as pre-computed in the Bgee database (‘gold’ set, version 14, (Bastian et al. 2008)). We studied the following tissues: lung, blood, placenta, liver, heart, brain, kidney, cerebellum, breast and testis. Importantly, most microRNA and mRNA extractions were done in the same commercial samples by the same lab (Brawand et al. 2011; Meunier et al. 2013), which suggest that co-expressed microRNA/mRNA pairs were present in the same tissue. MicroRNAs with more than 50 RPM (reads per million) were considered moderately expressed, and those with over 500 RPM were labelled as highly expressed. For transcripts we did not make distinctions between different expression levels, as transcript levels are affected by the action of microRNAs (Nam et al. 2014). Co-expressed microRNA:gene pairs were those whose microRNA was moderately or highly expressed in a tissue (depending on the analysis described) and the transcript targeted is detected in the same tissue. For ‘blood’ and ‘liver’ tissues we perform additional analysis considering the tissue specific polyadenilation signals as reported in APADB version 2 (Müller et al. 2014). MicroRNAs were grouped into evolutionary conservation categories depending on the species spread (primates, mammals and animals) of the seed family in TargetScan 7.2 (Agarwal et al. 2015).

Fst values were retrieved from the 1,000 Genomes Selection Browser 1.0 (Pybus et al. 2014). In Figure 4 the fold enrichment was computed as the logarithm of the ratio between the proportion of SNPs at target sites and the proportion of SNPs at a background site for each Fst bin. The latitude of specific populations was computed as in (Helmy et al. 2019). First, the geographical locations of the studied populations were obtained from the sample descriptions at https://www.coriell.org/1/NHGRI/Collections/1000-Genomes-Collections/1000-Genomes-Project., using the location of the capital of the country in cases where the location was ambiguous or nonspecific. For multiple collection points we computed the centroid middle point. Populations that migrated in recent times were discarded (CHD, CEU, ASW, ACB, GIH, STU and ITU as described at IGSR: The International Genome Sample Resource (http://www.internationalgenome.org/category/population/)).

All statistical analyses were done with R (v. 3.4.3, (R Development Core Team 2009)). All processed datasets and online tools to compute the tests here reported are available at our dedicated web server PopTargs (https://poptargs.essex.ac.uk; Hatlen and Marco, 2019). All the code use for the generation of the results, figures and tables is freely available from FigShare at https://doi.org/10.6084/m9.figshare.9539645.

## ACKNOWLEDGEMENTS

We thank Richard Nichols and Ben Skinner for critical reading of the manuscript and useful comments, and an editor and two anonymous reviewers for constructive criticisms on the methodology. We also acknowledge the use of the High Performance Computing Facility (Ceres) and its associated support services at the University of Essex in the completion of this work. This research was supported by the Wellcome Trust [grant number 200585/Z/16/Z].

## SUPPORTING INFORMATION LEGENDS

**Supplementary Figure 1.** Derived (target) allele frequency distributions for ten different tissues, compared with the expected distribution.

**Supplementary Figure 2.** Derived (target) allele frequency distributions for ten different tissues for highly expressed microRNAs, compared with the distribution for microRNAs with no detectable expression level.

**Supplementary Figure 3.** Derived (target) allele frequency distributions for wobble and non-wobble non-targets.

**Supplementary Table 1.** Statistical comparison of the derived (target) allele distributions at potential target sites versus background (expected) for 10 different tissues.

**Supplementary Table 2.** Statistical comparison of the derived (target) allele distributions at potential target sites of highly-expressed microRNAs versus target sites for non-detected (zero expressed) microRNAs for 10 different tissues.

**Supplementary Table 3.** P-values for the statistical support of effect of dependent variables and interaction in two linear models (see main text).

