## Supplementary figures and images for "Pervasive selection against microRNA target sites in human populations"

### Supplementary File 1

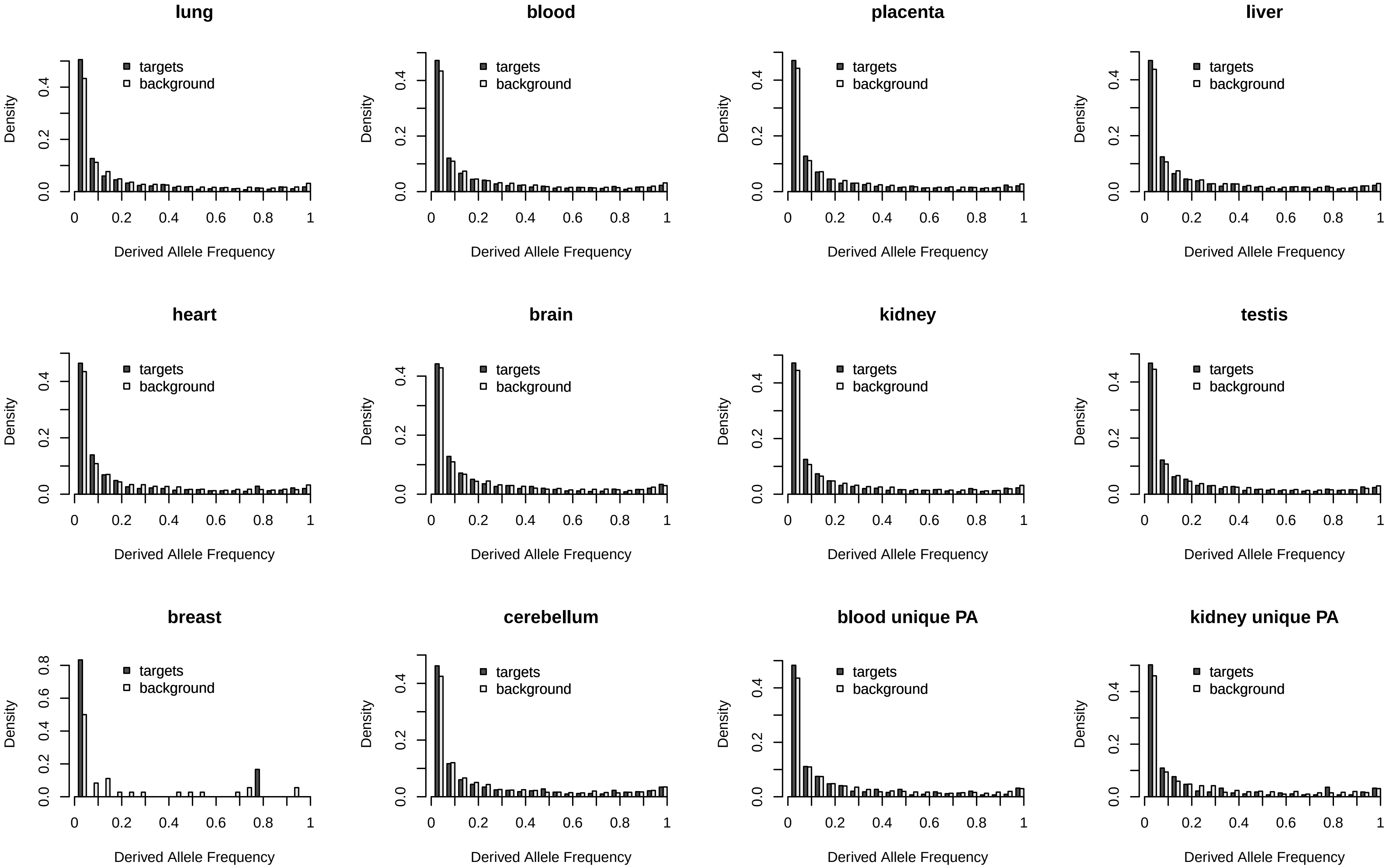

### Supplementary File 2

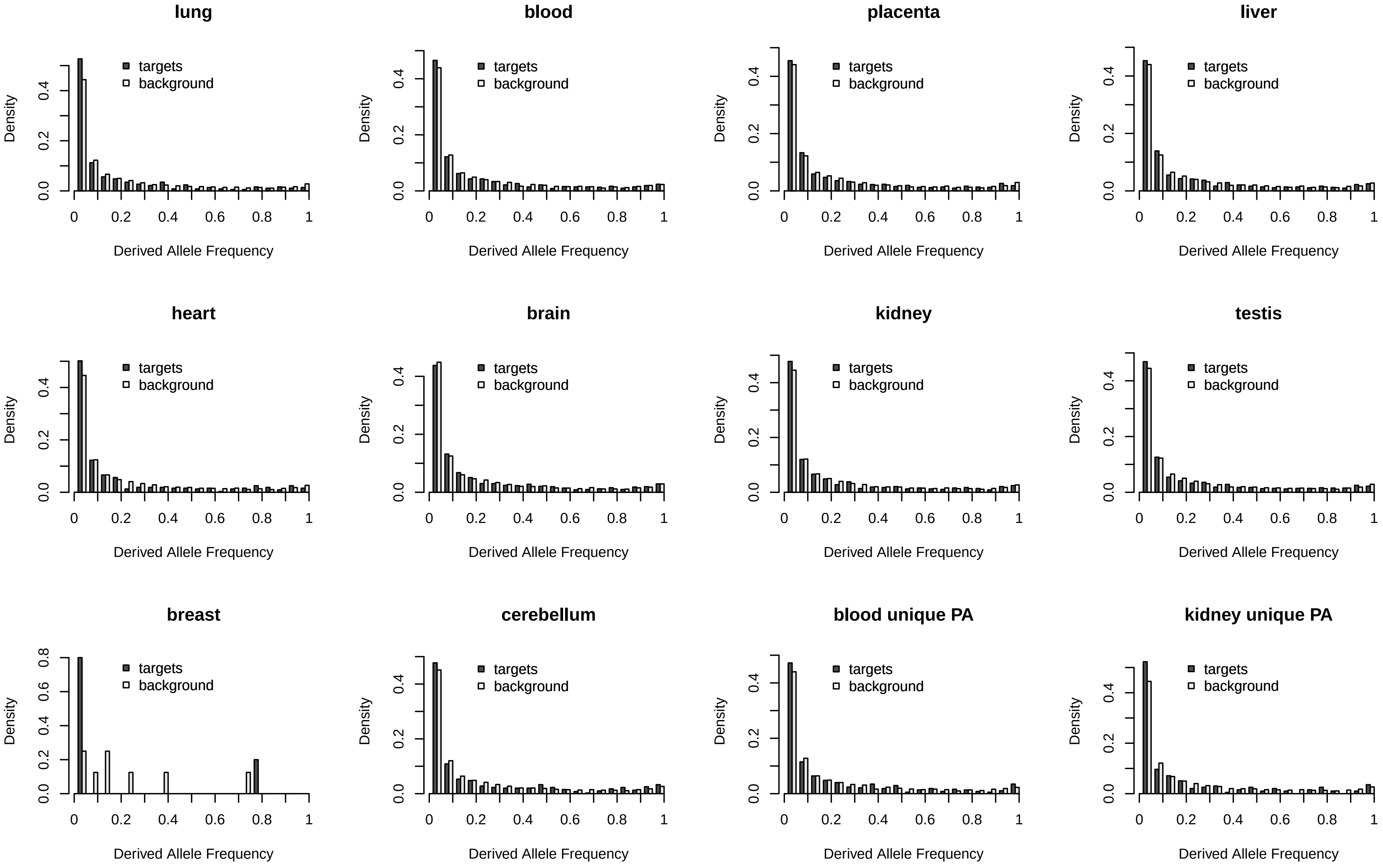

### Supplementary File 3

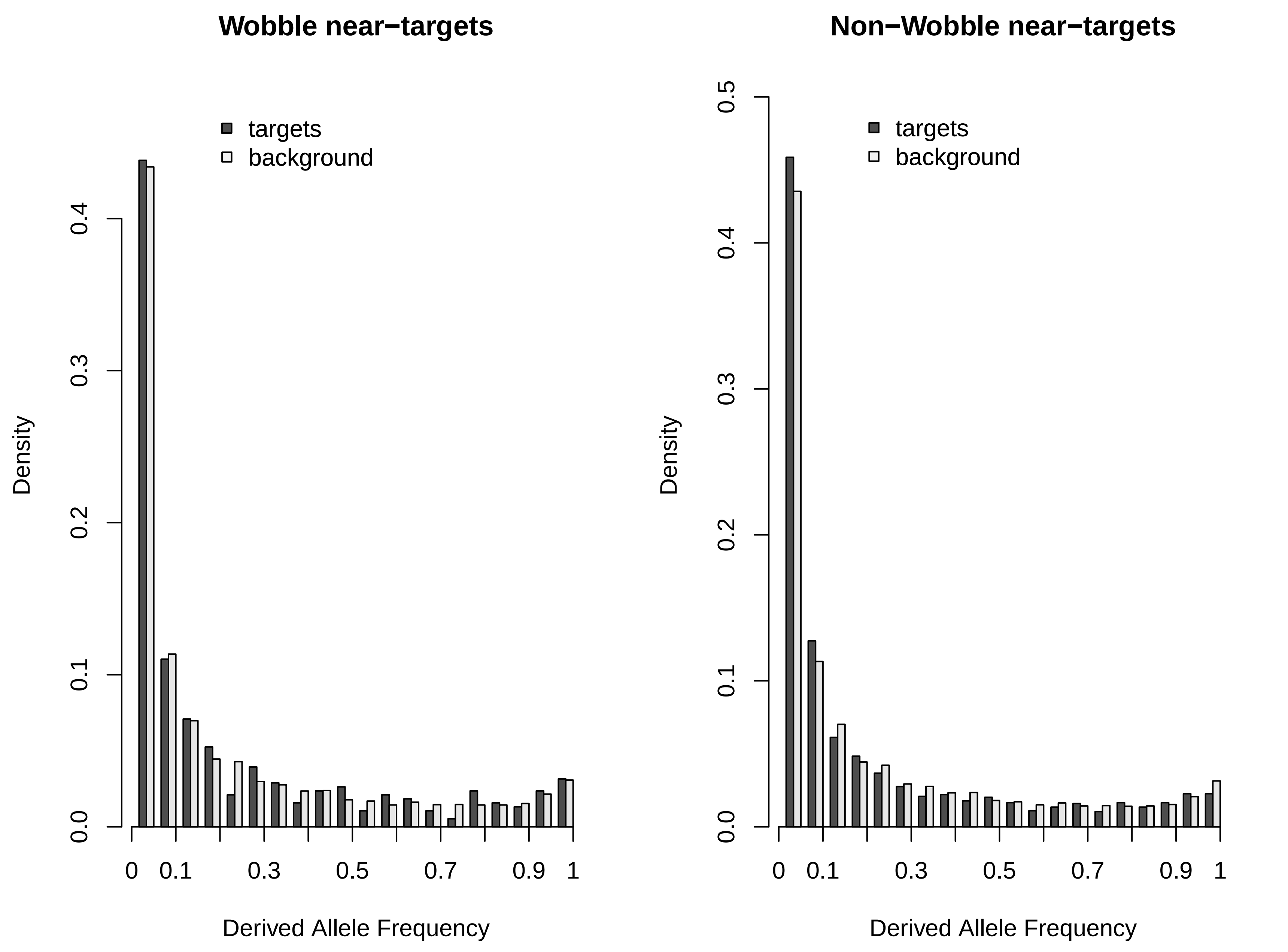
