## Supplementary Tables for "Pervasive selection against microRNA target sites in human populations"

**Supplementary Table 1.** Statistical comparison of the derived (target) allele distributions at potential target sites versus background (expected) for 10 different tissues.

| tissue | # SNPs interactions | # SNPs background | p[1] | q[2] |
|---|---|---|---|---|
| lung | 552 | 2292 | 0.000 | 0.001 |
| blood | 970 | 3907 | 0.003 | 0.016 |
| placenta | 1154 | 4644 | 0.006 | 0.018 |
| liver | 932 | 3803 | 0.007 | 0.018 |
| heart | 495 | 1927 | 0.017 | 0.028 |
| brain | 992 | 4232 | 0.026 | 0.037 |
| kidney | 791 | 3112 | 0.010 | 0.021 |
| testis | 1126 | 4552 | 0.038 | 0.045 |
| breast | 6 | 36 | 0.211 | 0.211 |
| cerebellum | 617 | 2339 | 0.040 | 0.045 |
| blood (unique PA) | 441 | 1872 | 0.027 | 0.034 |
| kidney (unique PA) | 275 | 998 | 0.034 | 0.034 |

[1] p-value computed from a one-tailed Kolmogorov-Smirnov test. [2] q-value (False Discovery Rate).

**Supplementary Table 2.** Statistical comparison of the derived (target) allele distributions at potential target sites of highly-expressed microRNAs versus target sites for non-detected (zero expressed) microRNAs for 10 different tissues.

| tissue | # SNPs interactions | # SNPs background | p[1] | q[2] |
|---|---|---|---|---|
| lung | 374 | 6838 | 0.001 | 0.005 |
| blood | 840 | 2630 | 0.027 | 0.068 |
| placenta | 1036 | 4177 | 0.196 | 0.245 |
| liver | 726 | 5488 | 0.113 | 0.189 |
| heart | 319 | 7421 | 0.005 | 0.023 |
| brain | 827 | 4189 | 0.688 | 0.688 |
| kidney | 576 | 7263 | 0.025 | 0.068 |
| testis | 922 | 6133 | 0.133 | 0.190 |
| breast | 5 | 8 | 0.061 | 0.121 |
| cerebellum | 396 | 7602 | 0.307 | 0.341 |
| blood (unique PA) | 375 | 2737 | 0.187 | 0.187 |
| kidney (unique PA) | 197 | 7455 | 0.034 | 0.068 |

[1] p-value computed from a one-tailed Kolmogorov-Smirnov test. [2] q-value (False Discovery Rate).

**Supplementary Table 3.** P-values for the statistical support of effect of dependent variables and interaction in two linear models (see main text). P-values below 0.05 are in red.

| | **Linear model (p-values)** | | |
|---|---|---|---|
| **Tissue** | **conservation** | **expression** | **interaction** |
| lung | 0.427 | 0.000 | 0.334 |
| blood | 0.163 | 0.156 | 0.262 |
| placenta | 0.165 | 0.182 | 0.295 |
| liver | 0.316 | 0.038 | 0.372 |
| heart | 0.940 | 0.020 | 0.393 |
| brain | 0.551 | 0.640 | 0.134 |
| kidney | 0.694 | 0.015 | 0.789 |
| testis | 0.945 | 0.055 | 0.361 |
| breast | 0.830 | 0.049 | 0.570 |
| cerebellum | 0.900 | 0.794 | 0.650 |
